# CysNet: Theorem constrained inference of cysteine redox proteoform states from bottom-up mass spectrometry data

**DOI:** 10.64898/2026.07.06.736853

**Authors:** James N. Cobley, Hao Jiang, Melpomeni Platani, Angus I. Lamond

## Abstract

Here, we present CysNet, a theorem-constrained method designed to infer cysteine redox proteoforms, i.e.,’oxiforms’, from bottom-up, mass spectrometry (MS)-based proteomic data. This overcomes limitations with previous MS redox proteomic approaches, which can quantify residue-resolved cysteine redox states, but leave distinct oxiforms unresolved. CysNet treats each residue-resolved oxidation value as a binary redox-coordinate marginal, enabling theorem-constrained inference of the oxiforms that are necessary, impossible or bounded within the compatible protein-group ensemble. This collapses the vast theoretically possible set of oxiform states to a finite set of allowed values by extracting existence and exclusion constraints from the data, despite the incomplete proteome coverage typical for bottom-up MS datasets. Using CysNet to analyse human induced pluripotent stem cell lines (~6,300 cysteine-containing protein groups, ~22% cysteine coverage), resolved 519 exact oxiforms, inferring 7,000 oxiforms per line. Quantitatively, CysNet bounded the oxiform content to 6.36–8.24 × 10^12^ protein copies, corresponding to 14–19% of the measured cysteine proteome. These data define the deepest oxiform survey recorded. CysNet revealed a latent structural layer in redox variation between the cell lines, distinguishing changes in oxiform identity (composition) from changes in oxiform weighting (intensity). Hence, CysNet moves bottom-up redox proteomics beyond isolated site-level cataloguing by reconstructing copy-number-weighted oxiform maps, providing a scalable route to deep oxiform information from peptide-level data.

## Introduction

Cysteine oxidation represents an important and widespread set of post-translational protein modifications that can affect protein structure and function. Cysteine oxidation provides a regulatory mechanism that acts via modulating the redox state of cysteine-containing protein sequences^1–3^. Even though cysteine oxidation is almost exclusively considered in isolation on a site-by-site basis^4^, redox regulation is encoded by combination of reduced and oxidised cysteine residues on proteoform molecules. These combinations define distinct cysteine proteoforms—oxiforms—which can differ in their resultant structural, catalytic, binding or allosteric properties^5–10^. That is, the structural and functional properties of the same protein molecule with one compared to two oxidised cysteines could differ, with implications for the biological logic of redox regulation.

Oxiform analysis is often assumed to require intact-protein, or top-down^11–13^, MS because the relevant object is the protein molecule carrying a specific combination of reduced and oxidised cysteine residues. However, intact-protein MS does not provide a simple direct readout of oxiform identity in complex biological extracts. The conceptual advantage of preserving molecular co-occupancy information is offset by the analytical difficulty of resolving small mass differences on larger and more heterogeneous analytes, where isotope envelopes, charge-state distributions, adducts, sequence variants and unrelated proteoforms can overlap^14,15^. Moreover, MS^2^ and MS^3^ necessarily convert the intact molecule into fragments, meaning that oxiform assignment still depends on inference from fragment-level information. Hence, meaningful oxiform assignment remains an inferential problem, and this inference has not been widely addressed in redox biology^16^. As a result, it remains unclear whether biological differences in cysteine redox state reflect changes in oxiform composition, where different oxiforms are present, or changes in oxiform weighting, where shared oxiforms differ in abundance.

Bottom-up MS presents the opposite problem. By digesting proteins into smaller, more regular peptide fragments before analysis, bottom-up workflows provide sensitive and scalable residue-level measurements. However, they are traditionally considered unable to recover protein-level oxiform information because proteolysis destroys direct molecular connectivity between cysteine sites^17,18^. The cost of this shorter fragment scale is loss of relational information: bottom-up data do not directly reveal which cysteine redox states, or other PTMs, co-occur on the same protein molecule. We reasoned, however, that if oxiform assignment is fundamentally an inferential problem even for intact-protein MS, then bottom-up redox proteomics might also support valid inference if the underlying oxiform space is given an explicit mathematical structure^19–21^. Specifically, by representing each protein group as a binary cysteine-redox state space, each residue-resolved cysteine oxidation value can be treated as a marginal constraint on an underlying oxiform distribution. In a structured state space, bottom-up data can determine which oxiform states are necessary, impossible or bounded by the observed peptide-level site information.

Here, we present CysNet, a theorem-constrained computational method provided as an open-source Python workflow that allows recovery of oxiform information from bottom-up MS peptide redox data. CysNet combines as input site-resolved cysteine redox measurements, FASTA-derived cysteine coordinates and protein-group quantities. For each protein group, cysteine residues are treated as binary redox coordinates and each measured cysteine oxidation value as a linear constraint on the distribution of compatible oxiform substates. CysNet reports which oxiform substates are compatible or excluded from the data, along with oxiform-existence constraints, feasible-state compression, resolution status and copy-number-scaled substate bounds. As a proof of principle, we used CysNet to analyse oxiform expression across eight unique human induced pluripotent stem cell lines (HIPSCI) derived from different donors. This revealed line-dependent differences in HIPSCI cysteine redox proteomes, detected as protein copy-number-scaled differences in thousands of oxiforms. CysNet demonstrates the ability to infer theorem-constrained proteoform-level information using high throughput, bottom-up MS redox proteomic data.

## Results

### Design of CysNet

CysNet was designed as a post-search, theorem-constrained computational workflow for converting site-resolved cysteine redox measurements into constraints on the oxiform ensembles compatible with bottom-up proteomic data (**Figure 1**). The workflow uses three inputs: site-level cysteine oxidation values, cysteine positions parsed from a FASTA file, and protein-group quantities.

**Figure 1.**
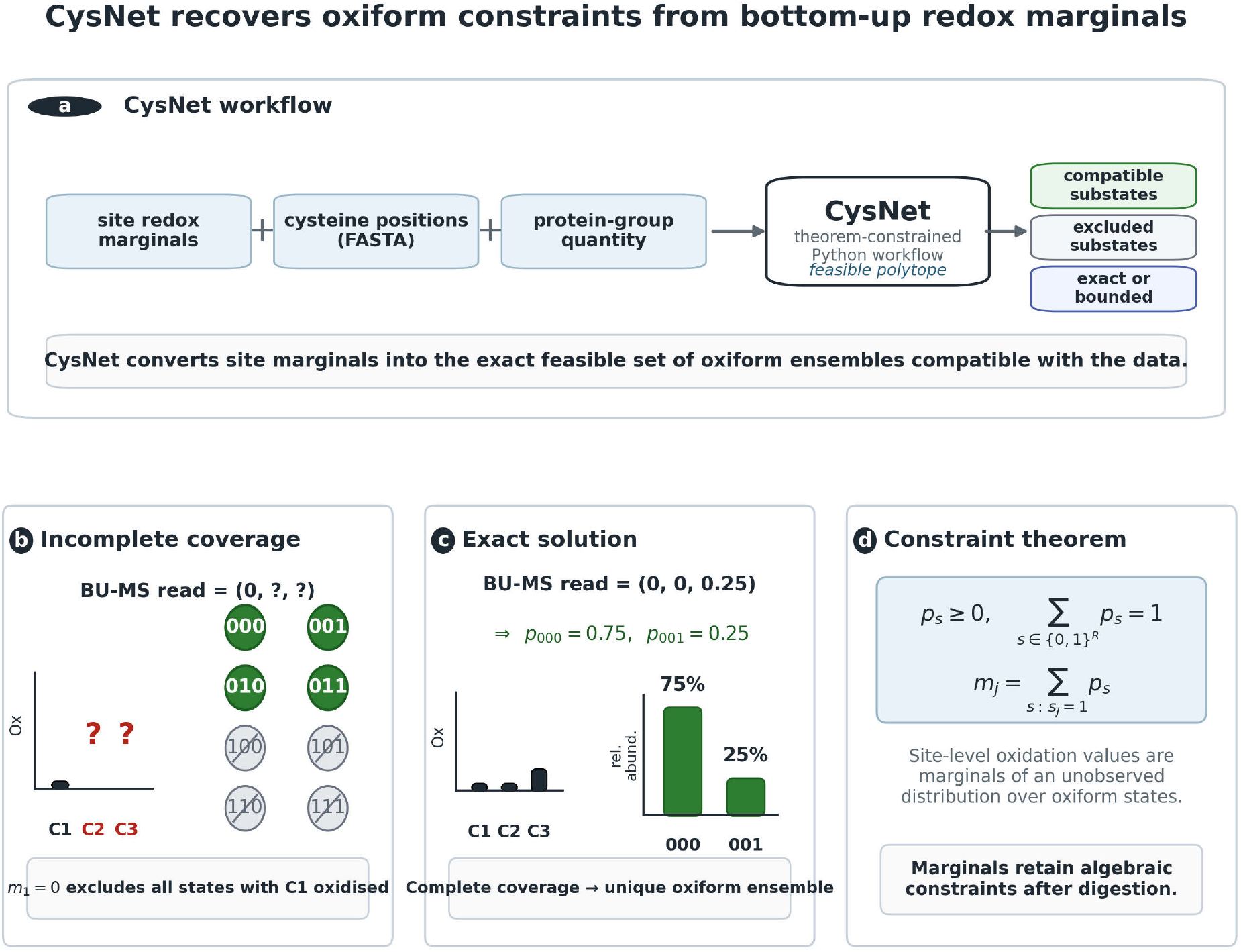
CysNet converts site-level redox marginals into constraints on compatible oxiform ensembles. **(A)**CysNet takes site-resolved cysteine redox marginals, FASTA-derived cysteine topology and protein-group quantities as input, constructs the feasible oxiform ensemble polytope and reports compatible, excluded, exact or bounded substates with copy-number-scaled bounds. **(B)** When cysteine coverage is incomplete, a measured boundary marginal excludes incompatible substates. **(C)** When cysteine coverage is complete, marginals can collapse the feasible set to an exact ensemble distribution. **(D)** The theoreom basis: site oxidation values are marginals over an unobserved binary oxiform distribution and therefore impose linear constraints after digestion.

The mathematical advance in CysNet is to give the oxiform problem an explicit combinatorial structure. For a protein group containing *R* measurable cysteine residues, each cysteine is represented as a binary redox coordinate that can occupy either state 0, denoting the reduced or unmodified form, or state 1, denoting the oxidised or modified form. An oxiform is therefore a specific binary cysteine-redox state of an individual protein molecule. Partial oxidation values, such as 25% oxidation (i.e., 0.25), arise at the population level and reflect the relative occupancy of binary oxiform states.

The set of possible oxiforms forms a binomial state space with 2^*R*^vertices, distributed across oxidation grades *k* = 0, …, *R*, where each grade contains 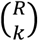 states. Geometrically, this state space can be represented as a graph in which each vertex is a possible oxiform state and edges connect states that differ by oxidation at a single cysteine coordinate. When organised into rows by oxidation grade *k*, this graph forms a diamond. The all-reduced state sits at *k* = 0, singly oxidised states at *k* = 1, higher-order oxiforms in successive rows, and the fully oxidised state at *k* = *R*. This provides a natural geometric representation of the state space, with cysteine sites acting as binary coordinates and oxiforms occupying vertices within the diamond.

The logic is illustrated by a protein containing three cysteines, corresponding to the modal cysteine count per protein in the human proteome. This protein has (2^3^ = 8) possible oxiform states in binary bitstring: 000, 001, 010, 011, 100, 101, 110 and 111. If the second cysteine is measured to be fully reduced, then every state in which this cysteine is oxidised is incompatible with the data. The states 010, 011, 110 and 111 are therefore excluded. A single boundary measurement has eliminated half of the theoretical state space and excludes the fully oxidised state 111.

If the first two cysteines are both measured to be fully reduced, the compatible state space collapses further to only 000 and 001. If the measured oxidation values for the three cysteines are 0, 0 and 0.25, then the compatible oxiform ensemble is uniquely determined: 75% of molecules occupy state 000 and 25% occupy state 001. This example shows how site-level measurements from bottom-up MS can resolve a protein-level oxiform ensemble when the measured residue-resolved redox values uniquely constrain the underlying binary state distribution.

CysNet generalises this logic across protein groups by constructing the oxiform state space for each protein group and applying the experimentally measured residue-resolved cysteine oxidation values as linear marginal constraints. At its core, CysNet implements a theorem on this mathematical structure, which defines what can be inferred from the measured marginals: which oxiforms are compatible, excluded, necessary or bounded, and what abundance constraints follow from the data (**Supplemental information**). The resulting protein-level output includes mathematically constrained oxiform-level information from the bottom-up redox proteomic data, such as copy-number-scaled oxiform abundance. This moves bottom-up proteomics from considering cysteine residues in isolation toward quantitative oxiform mapping.

### Oxiform inference from hypothetical bottom-up redox proteomic data

To illustrate the mathematical object recovered by CysNet, we considered a synthetic redox proteomic data set. For a protein group with *R* cysteines, each molecule in the population must adopt one oxiform state. These molecular states can be represented as a binary vector

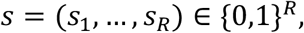

where *s*_*j*_ = 0 denotes a site with a reduced cysteine and *s*_*j*_ = 1 denotes a site with an oxidised cysteine at the single molecule state level. The full oxiform state space is therefore

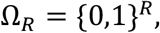

with

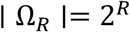

possible oxiform states^6^. These oxiform states can also be grouped by oxidation grade,

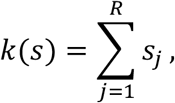

where *k* is the number of oxidised cysteines in the state. For *R* = 3, the state space contains eight substates distributed across four oxidation grades per the binomial theorem^20^

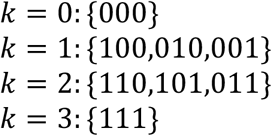

More generally, grade *k* contains

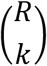

states. Hence, before any redox measurement is made, the protein group has a theoretical combinatorial oxiform state space, whose size grows exponentially with cysteine number^8^.

An unresolved oxiform ensemble is a distribution over this entire theoretical state space. We write *p*_*s*_ for the fractional abundance of substate *s*, with

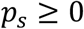

and

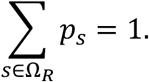

Note that a measured redox value for a single cysteine site defines a parameter (termed a ‘site marginal’) related to the hidden distribution of actual oxiforms that exist in the sample. For cysteine *j*, the measured oxidation value *m*_*j*_ satisfies

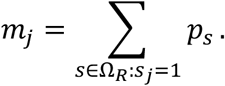

Each residue-resolved cysteine redox value measured in a bottom-up MS data set provides a linear constraint on the unknown distribution of actual oxiform states that exist in the sample. If only a subset *J* ⊆ {1, …, *R*} of the total number of cysteines is measured, then the feasible oxiform ensemble is the mathematical solution to the diamond-shaped graph (i.e., the convex polytope)

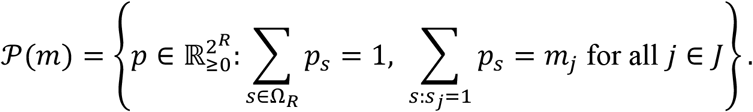

CysNet infers this feasible set (**Figure 2A**). Depending on the information content of the measured cysteine marginals, the feasible set of actual oxiforms present in a sample may remain broad, become bounded, exclude specific substates, require the presence of oxidised substates, or collapse to a single, exact ensemble distribution.

**Figure 2.**
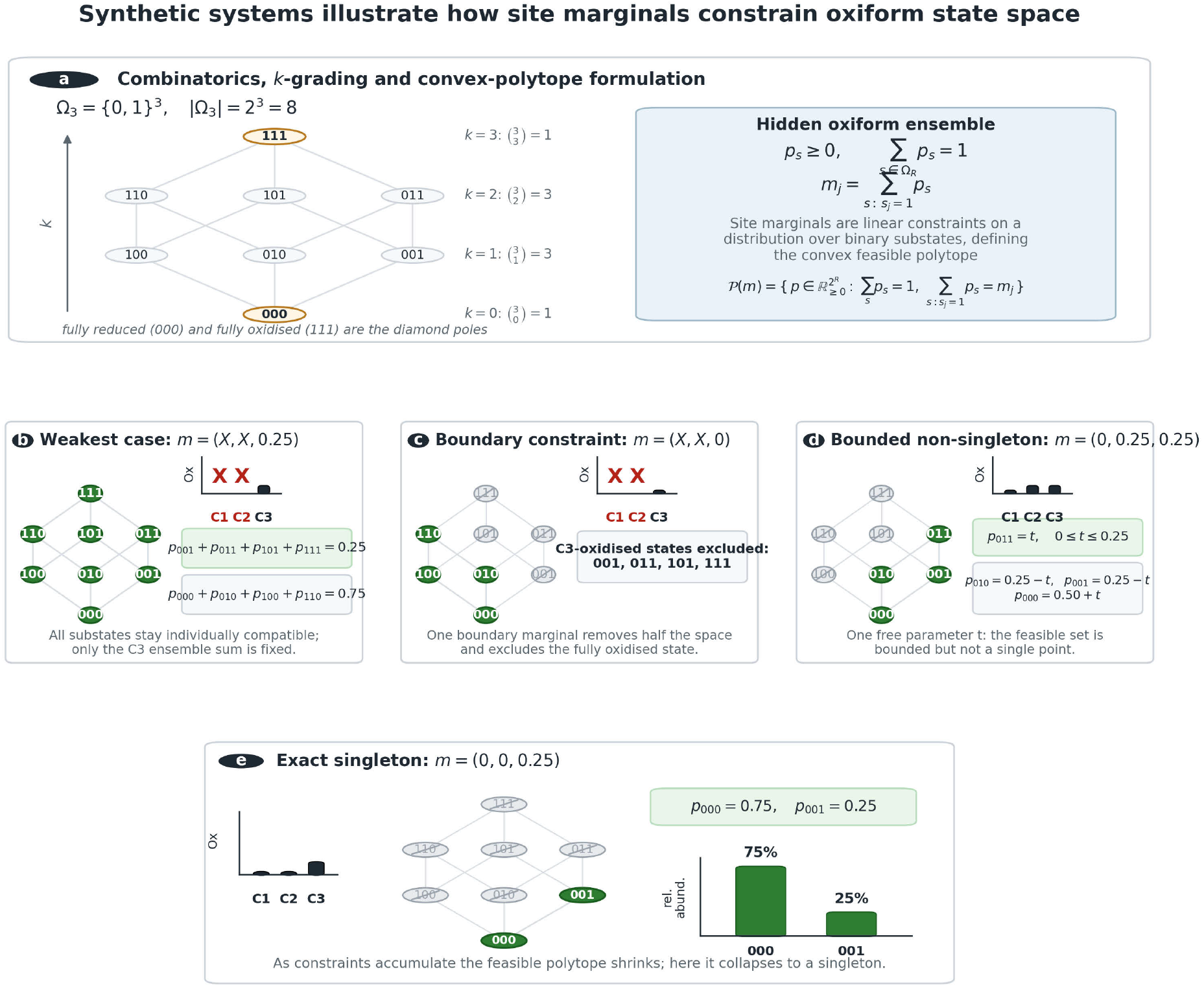
Oxiform inference from hypothetical redox proteomic data. **(A)** For a protein with *R* = 3cysteines, the possible cysteine-redox substates form the binary state space Ω_3_ = {0,1}^3^, containing | Ω_3_ |= 2^3^ = 8 substates. These substates can be graded by oxidation number *k*, where *k* = 0corresponds to the fully reduced state 000, *k* = 3 to the fully oxidised state 111, and intermediate grades contain 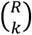 substates. An unresolved oxiform ensemble is a distribution *p*_*s*_over this state space, with *p*_*s*_ ≥ 0 and ∑_*s*_ *p*_*s*_ = 1. Each measured site oxidation value *m*_*j*_is a marginal of this hidden distribution,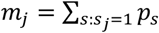, and therefore defines a linear constraint on the convex feasible polytope of compatible oxiform ensembles. **(B)** In the weakest illustrated case, only C3 is measured and is partially oxidised, *m* = (*X, X*, 0.25), where *X* denotes an unmeasured coordinate. No individual substate is excluded, but the ensemble is constrained: the total probability assigned to C3-oxidised substates must equal 0.25, *p*_001_ + *p*_011_ + *p*_101_ + *p*_111_ = 0.25, while the total probability assigned to C3-reduced substates must equal 0.75, *p*_000_ + *p*_010_ + *p*_100_ + *p*_110_ = 0.75. **(C)** A boundary marginal imposes a stronger constraint. When *m* = (*X, X*, 0), every C3-oxidised substate is incompatible with the data. CysNet therefore excludes 001, 011, 101, 111, including the fully oxidised state 111. **(D)** With *m* = (0,0.25,0.25), the C1 boundary marginal first excludes all C1-oxidised substates, while the two partial marginals constrain the remaining compatible substates without yielding a unique solution. The feasible set is a bounded non-singleton polytope parameterised by *p*_011_ = *t*, with 0 ≤ *t* ≤ 0.25. The remaining compatible substate abundances are then *p*_010_ = 0.25 − *t, p*_001_ = 0.25 − *t*,and *p*_000_ = 0.50 + *t*. Hence, the data bound the compatible oxiform ensemble but do not collapse it to a single point. **(E)** With *m* = (0,0,0.25), boundary constraints at C1 and C2 leave only 000and 001compatible with the data. The C3 marginal then fixes the unique ensemble distribution, *p*_000_ = 0.75 and *p*_001_ = 0.25.

To illustrate this progression, we considered a hypothetical protein with three cysteines. In the weakest case, only the redox state of the third cysteine is measured, and this cysteine is seen to be partially oxidised:

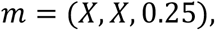

where *X* denotes an unmeasured residue (i.e., coordinate on the diamond-shaped graph). In this case, no oxiform state is excluded, because C1 and C2 are unconstrained and C3 is neither fixed as fully reduced nor fully oxidised (**Figure 2B**). However, the ensemble (population of molecules) is still constrained. The total abundance of all substates with C3 oxidised must equal 0.25:

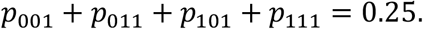

Likewise, the total abundance of all substates with C3 reduced must equal 0.75:

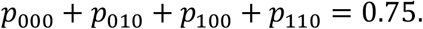

All eight potential oxiform substates remain individually compatible, but the ensemble cannot distribute probability mass arbitrarily. For example, the fully oxidised substate 111 remains possible, but it cannot occupy more than 25% of the ensemble because it contributes to the C3-oxidised marginal.

A boundary marginal imposes a stronger constraint. If the third cysteine is observed as fully reduced,

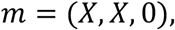

then every theoretical substate in which C3 is oxidised is incompatible with the experimental MS data (**Figure 2C**). The excluded states are

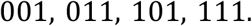

The state space compatible with the experimentally measured data is reduced from eight states to four:

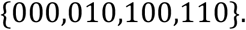

This shows that a single boundary marginal can remove half of the theoretical binary oxiform state space and exclude the existence of any molecules in the fully oxidised state. Constraints on the allowed oxiforms that can exist in the sample increase as more cysteine are measured. For example, consider

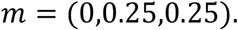

The boundary marginal at C1 excludes every potential substate in which C1 is oxidised, leaving only

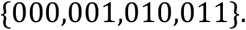

The two partial marginals then impose abundance constraints over the remaining compatible substates:

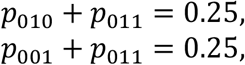

and

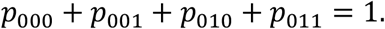

This system is constrained, but inexact (**Figure 2D**). If we let

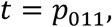

then the feasible ensemble can be written as

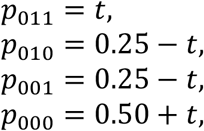

with

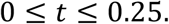

The data bound every compatible state without forcing a single ensemble distribution:

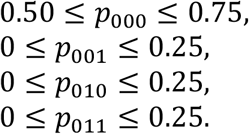

This bounded, non-singleton case shows that CysNet cannot over-resolve proteoform distributions beyond the information that is present in the MS data (i.e., it cannot by formal construction “hallucinate” or overclaim more than the data allow).

Finally, if two boundary marginals and one partial marginal are observed,.

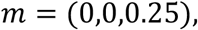

then C1 and C2 are both fixed as reduced. All states containing oxidation at either coordinate are excluded, leaving only

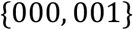

compatible with the data. The C3 marginal then fixes the unique compatible ensemble:

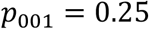

and

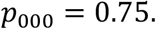

In this case, the feasible polytope collapses to a singleton and CysNet infers the exact oxiform ensemble distribution, with a quantitative weight (i.e., 000 is more abundant than 001, **Figure 2E**).

These hypothetical examples show how every measured cysteine redox value contributes information about the hidden oxiform distribution. CysNet therefore converts bottom-up MS redox measurements into the precise mathematical object retained after digestion: the feasible oxiform state space, including the compatible states, excluded states and abundance bounds implied by the data.

### Human iPSC proteomes reveal biological redox variation

We next used CysNet to characterise and compare oxiform distributions across eight different human induced pluripotent stem cell (iPSC) lines in the HIPSCI cohort. Each line was derived from a genetically distinct female donor, providing a tractable model of interindividual variability^22,23^. To generate empirical redox data for CysNet, the cell lines were grown in the pluripotent state and extracts prepared and used in a bottom-up, redox proteomics MS workflow, using the <u>re</u>dox <u>cap</u>ture (ReCap^24^) technique (**see methods**) Leveraging the cysteine oxidation state encoded data-independent acquisition (Oxi-DIA) on the Orbitrap-Astral™ coupled to the DIA-NN analysis pipeline^25–29^, the resulting data showed cell line-dependent variability in cysteine redox states. This variation in cysteine redox values was a function of the cell line and not seen between technical replicates (**Figure 3A-B**). Using Oxi-DIA, approx. 20,000 cysteine residues were quantified per donor (**Figure 3C**). Most of the detected cysteine sites were reduced (**Figure 3D**). Of 11,964 cysteine sites measured in each donor cell line, 7,953 had identical redox states in every biological sample, whereas 4,011 varied^30^ (**Figure 3E**). Hence, conventional site-level analysis established a iPSC cell line-associated, cysteine-redox phenotype (**Figure 3F**).

**Figure 3.**
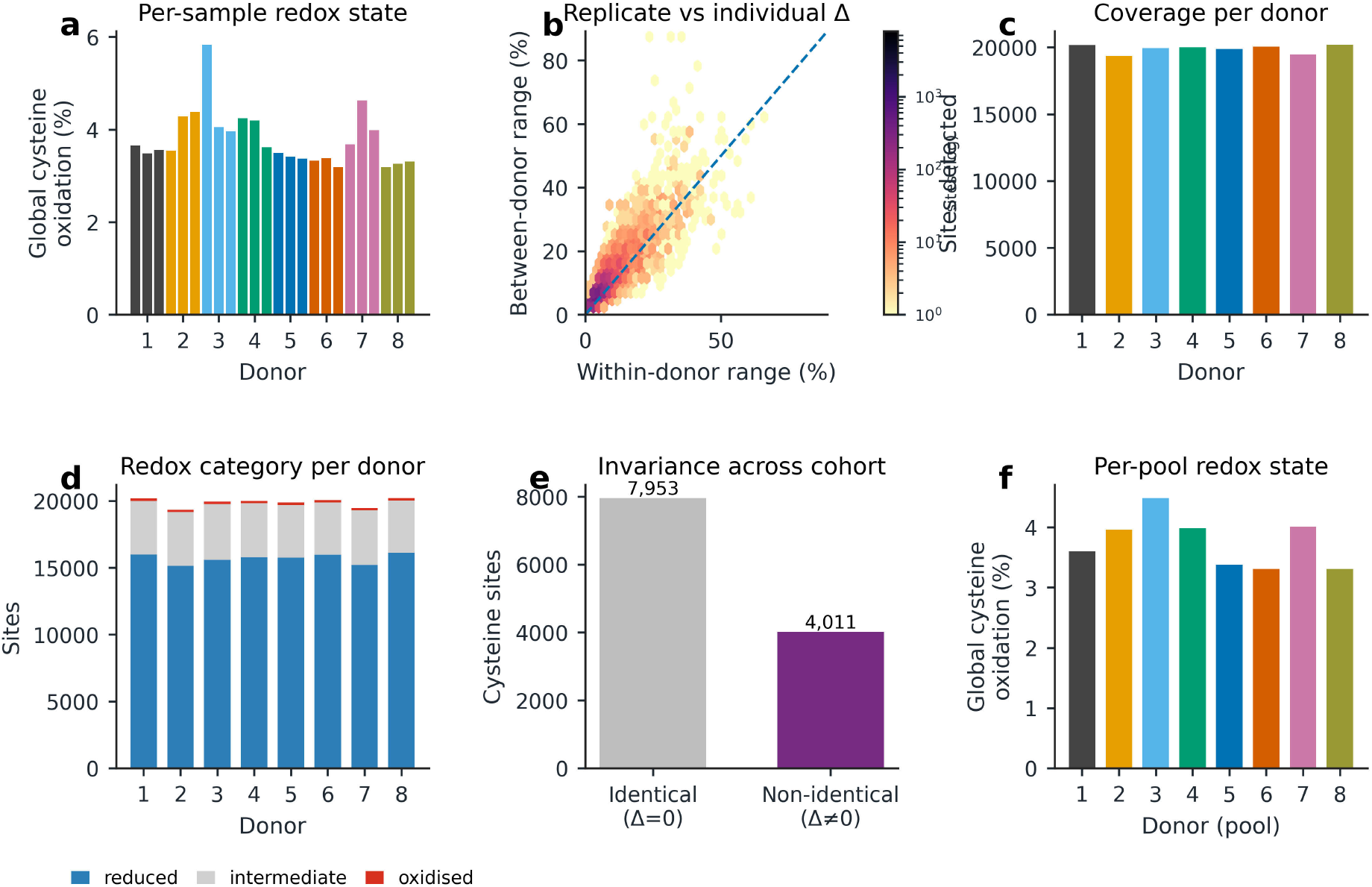
Biological cysteine-redox variation in human iPSC lines. **(A)** Global cysteine oxidation across individual bottom-up redox proteomics samples from eight unique female donor-derived human induced pluripotent stem cell lines. Bars represent the mean oxidation state of detected cysteine sites in each sample, with colours indicating donor identity. **(B)** Comparison of technical/within-donor variation and between-donor variation for cysteine sites detected across the full sample set. The x axis shows the mean within-donor range across replicate samples, whereas the y axis shows the range across donor-level means. The dashed line indicates equal within- and between-donor variation; sites above this line show greater donor-associated variation than replicate variation. **(C)** Number of detected cysteine sites per donor after donor-level aggregation. **(D)** Classification of detected cysteine sites per donor into fully reduced, intermediate and fully oxidised redox categories. Fully reduced sites had 0% oxidation, fully oxidised sites had 100% oxidation and intermediate sites had oxidation values between these boundaries. **(E)** Classification of cysteine sites detected across all samples as invariant or variable across the cohort. Identical (Δ= 0) sites had no observed redox difference across the full cohort, whereas non-identical sites (Δ≠ 0) showed at least one donor- or sample-associated redox difference. **(F)** Global cysteine oxidation after donor-level pooling, showing donor-dependent differences in the average redox state of the measured cysteine proteome.

Next, donor-level protein-group intensities were normalised to the injected protein mass and converted to approximate protein copy numbers, using FASTA-derived molecular weights, via the total protein approach^31^ (**Figure 4**). Weighting cysteine oxidation values by parent protein copy number yielded donor-specific, mean proteome-wide oxidation values ranging from 6.91% to 9.72%, with donors 3 and 7 showing the highest copy-weighted oxidation and donor 8 the lowest. Because site-level Oxi-DIA does not resolve whether oxidised cysteines co-occur on the same protein molecule, the number of molecules carrying at least one oxidised measured cysteine could only be bounded, not precisely determined.

**Figure 4.**
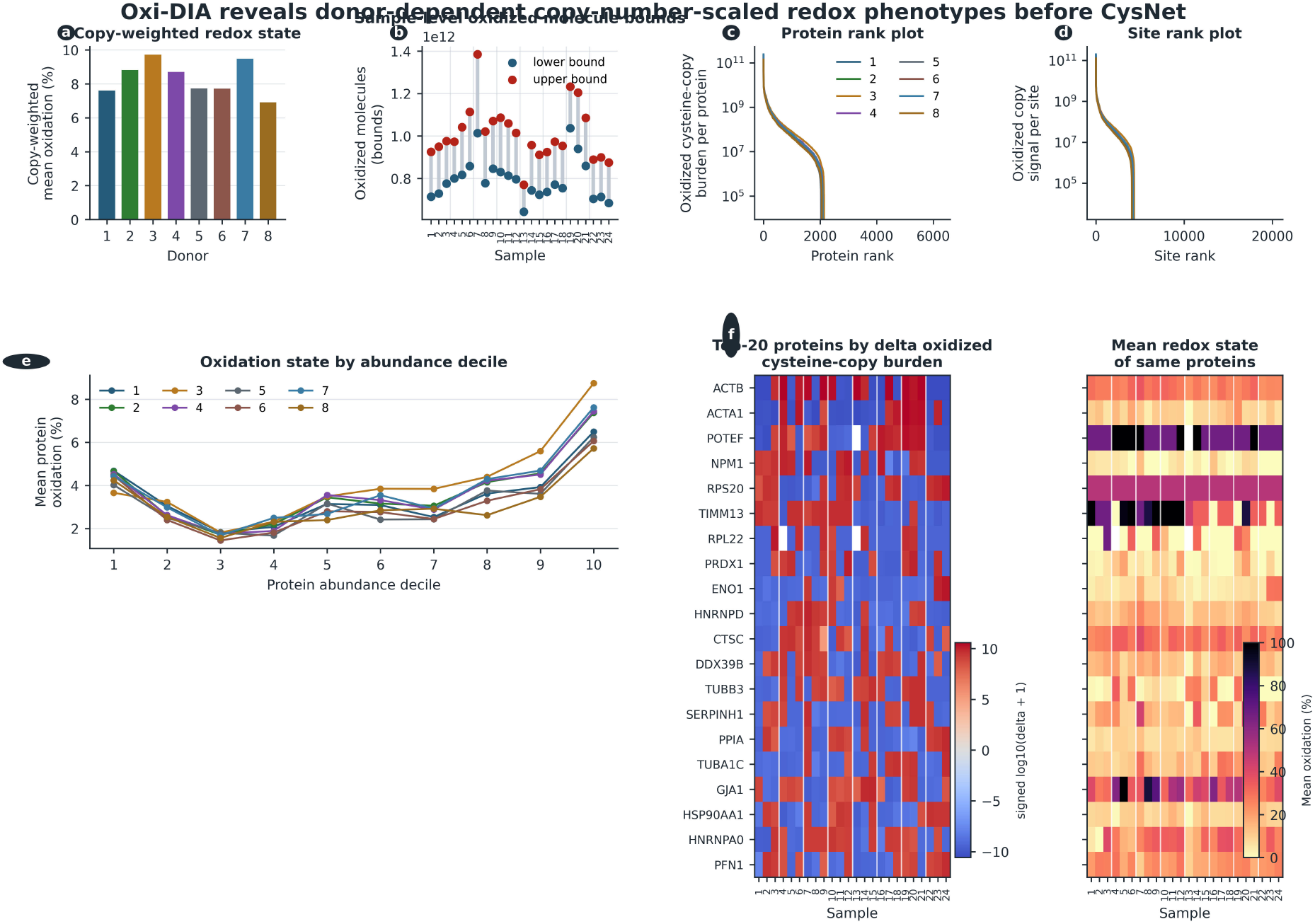
Copy-number scaling reveals donor-dependent Oxi-DIA redox burden before CysNet. **(A)** Copy-weighted mean cysteine oxidation per donor, calculated by weighting detected site oxidation values by the estimated copy number of the parent protein group. **(B)** Sample-level bounds on the number of protein molecules carrying at least one oxidised measured cysteine. Bounds were calculated per protein using the maximum detected site oxidation as a lower bound and the summed detected site oxidation, capped at one molecule, as an upper bound, then summed across proteins. These bounds reflect the measured cysteine coordinate space and do not resolve cysteine co-occurrence. **(C)** Rank plot of protein-level oxidised cysteine-copy burden, calculated as parent protein copy number multiplied by the sum of detected site oxidation fractions for each protein. **(D)** Rank plot of site-level oxidised copy signal, calculated as parent protein copy number multiplied by the oxidation fraction of each detected cysteine site. **(E)**, Mean protein oxidation across protein abundance deciles for each donor. **(F)** Top 20 proteins contributing most to sample-level variation in oxidised cysteine-copy burden, shown as signed log-transformed deviation from the sample mean, alongside the corresponding mean redox state of the same proteins.

Across samples, these bounds ranged from 6.43 × 10^11^ to 1.04 × 10^12^ molecules for the lower bound and from 7.70 × 10^11^ to 1.38 × 10^12^ molecules for the upper bound. Rank plots showed that oxidised copy signal was highly concentrated across both proteins and cysteine sites, By comparing how redox values varied in different protein groups according to their measured abundance class, this decile analysis showed a systematic relation between redox state and protein abundance The largest sample-level differences in oxidised cysteine copy numbers were driven by a small subset of highly abundant proteins, including ACTB, ACTA1, NPM1, RPS20, PRDX1, ENO1, HSP90AA1 and PPIA.

In sum, these data obtained using the Oxi-DIA MS method establishes donor-dependent variability in cysteine redox values, at both site and protein-copy scale.

### CysNet infers exact oxiforms from empirical iPSC bottom-up redox proteomic data

As described above, data derived using Oxi-DIA established line-associated variation in cysteine-redox states and copy-number scaling showed this represents a substantial difference in oxidation between the different donor cell lines. However, these analyses remain residue-centred, quantifying the relative redox status of individual cysteine sites and how much copy-number-weighted signal each site, or protein contributes. Next, therefore, we applied CysNet to analyse the same Oxi-DIA data to infer the oxiform-level constraint layer latent in empirical bottom-up redox data, allowing a comparison of oxiform ensembles between the respective hiPSC lines.

CysNet analysed 6,133–6,319 cysteine-containing protein groups and 19,337–20,195 detected cysteine coordinates per donor cell line, corresponding to approx. 22.1–22.5% coverage of total predicted cysteine residues in the proteomes expressed in these cell lines (**Figure 5A-C**). The theoretical cysteine proteoform state space belonging to these expressed protein groups was astronomically large, i.e., ~3.02 × 10^169^ in most donors. This was in part inflated by the presence of spondin (A2VEC9, **Figure 5D**), a protein with 563 cysteine residues, the most in the human proteome^8^. Spondin was not detected in donor eight. The protein with the most cysteine residues in donor eight Laminin subunit alpha 5 (*n* = 220, O15230). Even though the residue coverage was high (**Figure 5E**), complete cysteine coverage, i.e., detection of all the cysteine residues belonging to the protein group, was rare (**Figure 5F**).

**Figure 5.**
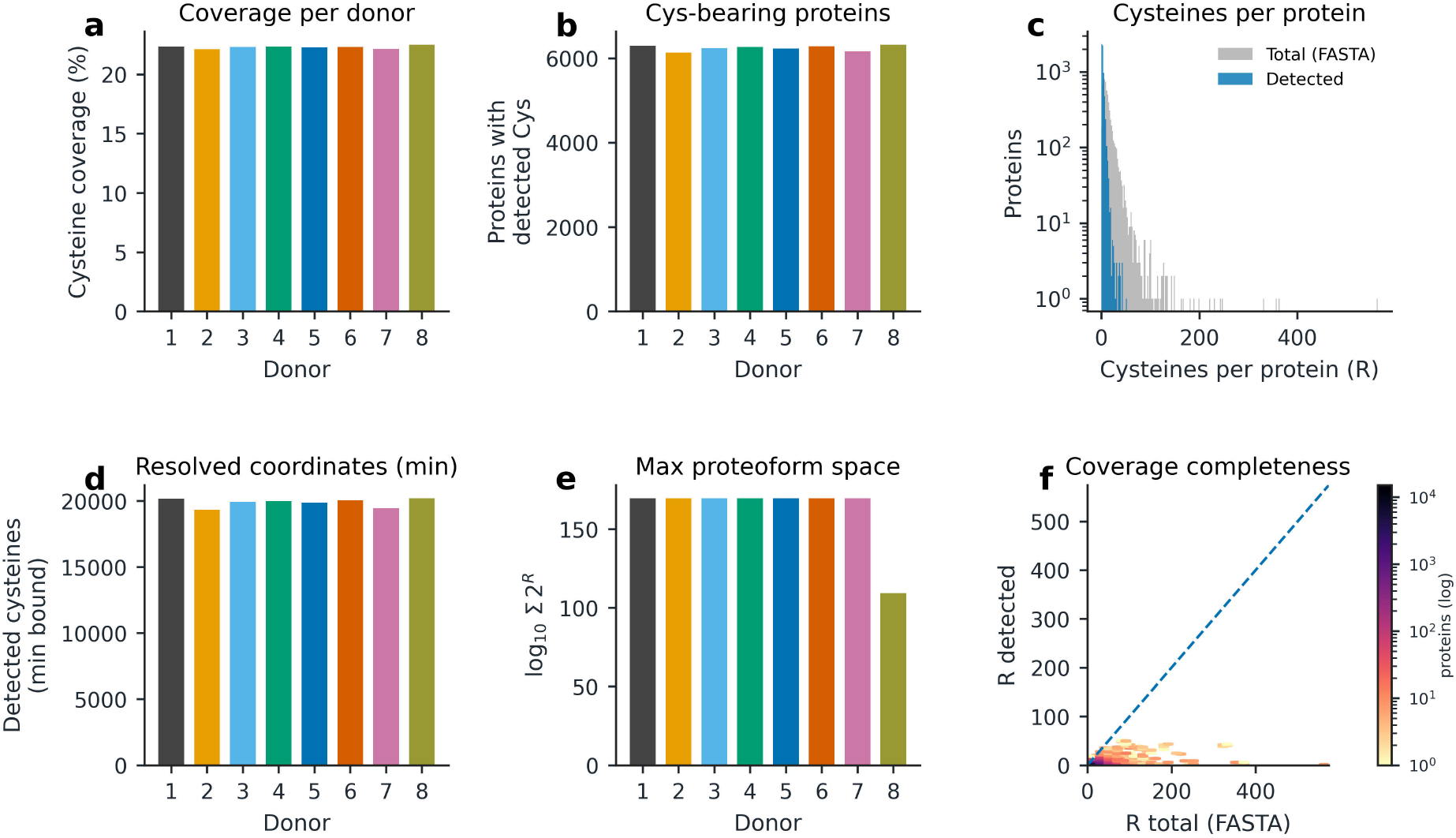
Cysteine coverage and theoretical oxiform state-space size across donor-derived iPSC redox proteomes. **(A)** Percentage cysteine coverage per donor, calculated as the fraction of FASTA-annotated cysteine residues detected by bottom-up redox proteomics. Coverage was consistent across the eight donor-derived iPSC lines, at approximately 22% of the cysteine-containing proteome. **(B)** Number of cysteine-bearing protein groups detected per donor. Each donor yielded approximately 6,000 cysteine-containing proteins with at least one detected cysteine coordinate. **(C)** Distribution of cysteine counts per protein in the reference FASTA compared with the subset of detected cysteine coordinates. Grey bars show total cysteine counts per protein in the reference proteome, whereas blue bars show detected cysteine counts per protein. The distribution illustrates the sparsity of bottom-up cysteine coverage relative to the full cysteine topology. **(D)** Minimum number of resolved cysteine coordinates across donors, showing the number of detected cysteine sites retained after applying donor-level detection criteria. **(E)** Maximum theoretical oxiform state-space size per donor, expressed as log_10_(2^*R*^), where *R*is the number of cysteine residues considered for each detected protein. This illustrates the combinatorial scale of the proteoform space implied by cysteine topology before CysNet constraint pruning. **(F)** Relationship between total FASTA cysteine count per protein and detected cysteine count per protein. The dashed line indicates complete cysteine coverage, where *R*_detected_ = *R*_total_. Most proteins fall below this line, confirming that empirical bottom-up redox data are dominated by incomplete cysteine coverage. Colour indicates the number of proteins on a logarithmic scale.

**Figure 6.**
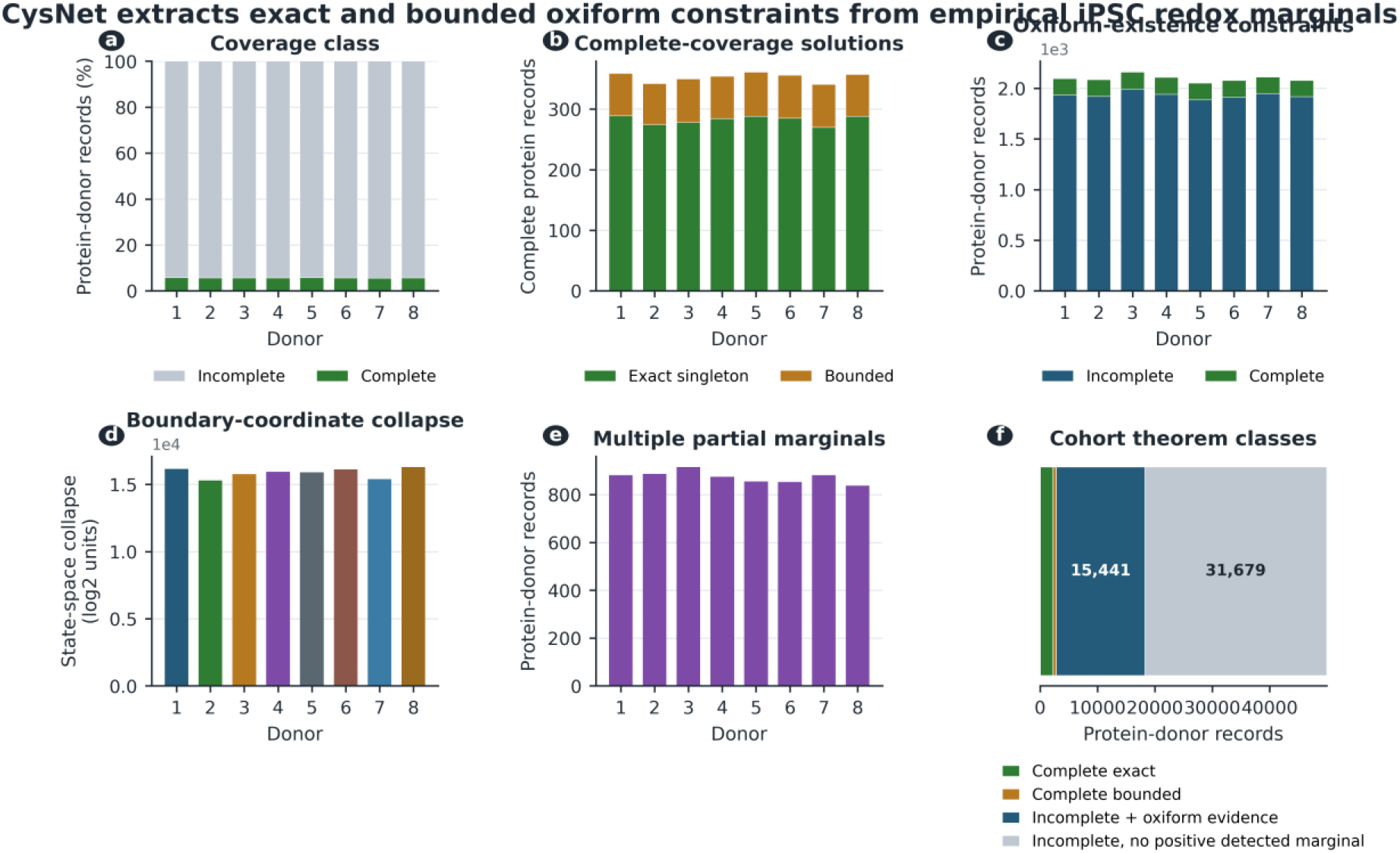
CysNet extracts exact and bounded oxiform constraints from empirical iPSC redox marginals. **(A)** Classification of protein-donor records according to cysteine coverage. Complete records contain all FASTA-annotated cysteine coordinates for the protein group, whereas incomplete records contain one or more missing cysteine coordinates. **(B)** Resolution status for complete-coverage records. Exact singleton denotes a feasible oxiform ensemble polytope that collapses to a single distribution; bounded denotes a non-singleton feasible polytope. **(C)** Protein-donor records with oxiform-existence evidence, defined as at least one positive measured cysteine oxidation marginal. In incomplete records, this proves that at least one oxidised substate is required within the detected coordinate space, without assigning a unique intact molecular state. **(D)**, Aggregate state-space collapse induced by boundary marginals. Each measured cysteine fixed at 0% or 100% removes one binary dimension from the compatible state space; values show the cumulative log2 collapse per donor. **(E)** Protein-donor records containing two or more intermediate cysteine marginals, representing cases in which partial redox coordinates impose bounded ensemble constraints. **(F)**, Cohort-level theorem classes across all protein-donor records. The figure separates complete exact solutions, complete bounded solutions, incomplete records with oxiform-existence evidence and incomplete records with no positive detected marginal. These classes summarise the constraint information retained in empirical bottom-up redox measurements before copy-number scaling.

**Figure 7.**
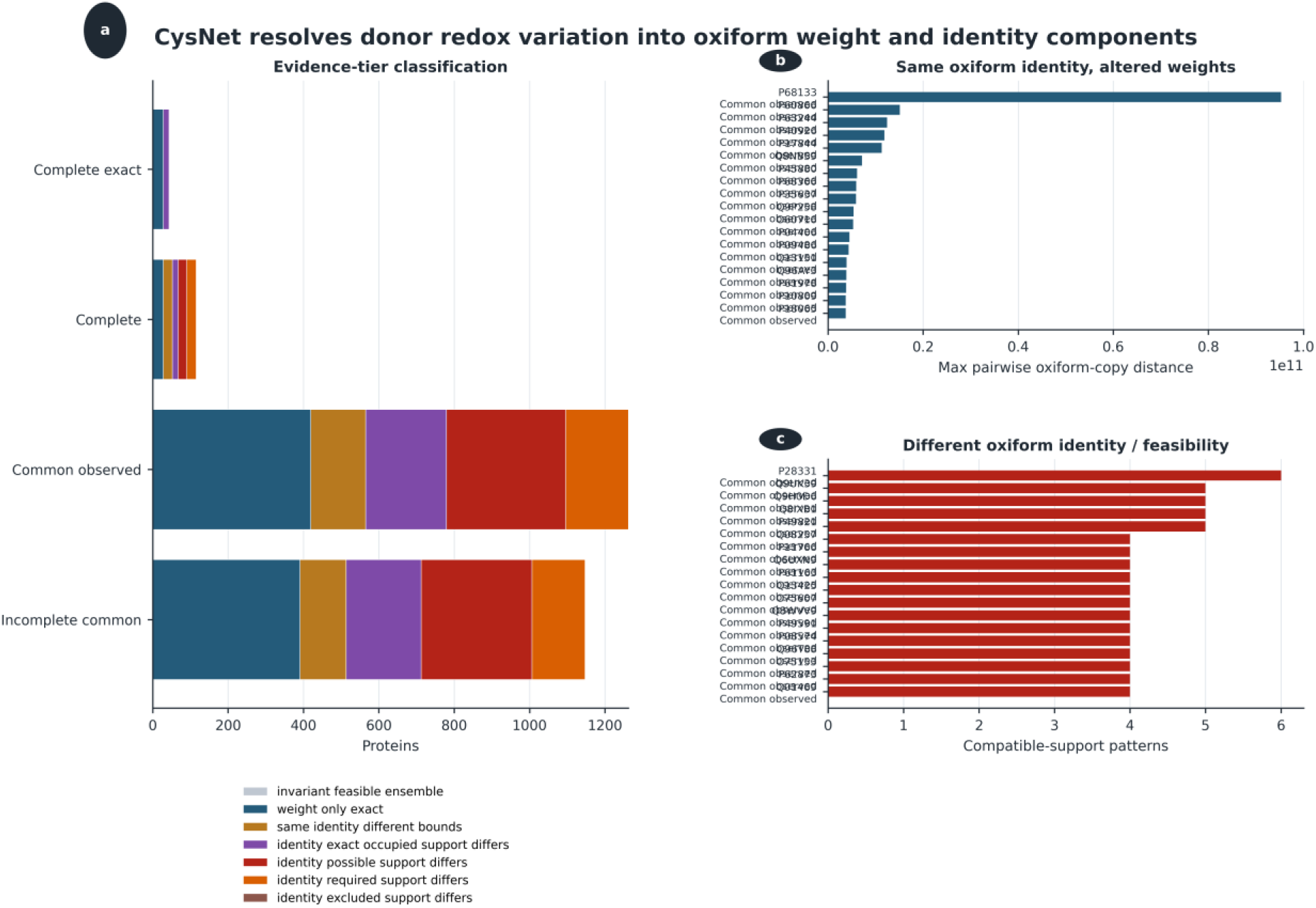
CysNet decomposes donor-dependent redox variation into oxiform identity and oxiform weighting. **(A)**, Evidence-tier classification of protein groups across donor comparisons. CysNet separates invariant feasible ensembles from cases in which donors share the same oxiform identity but differ in oxiform weight, and cases in which oxiform identity or feasibility differs through altered bounds, occupied support, possible support, required support or excluded support. **(B)** Protein groups with shared oxiform identity but altered weights, ranked by maximum pairwise oxiform-copy distance, identifying cases where the same compatible oxiform states are retained but their copy-number-scaled abundances differ between donors. **(C)** Protein groups with different oxiform identity or feasibility, ranked by the number of compatible-support patterns, identifying cases where donor variation reflects a change in the set of oxiform states permitted or supported by the measured cysteine-redox constraints.

**Figure 8.**
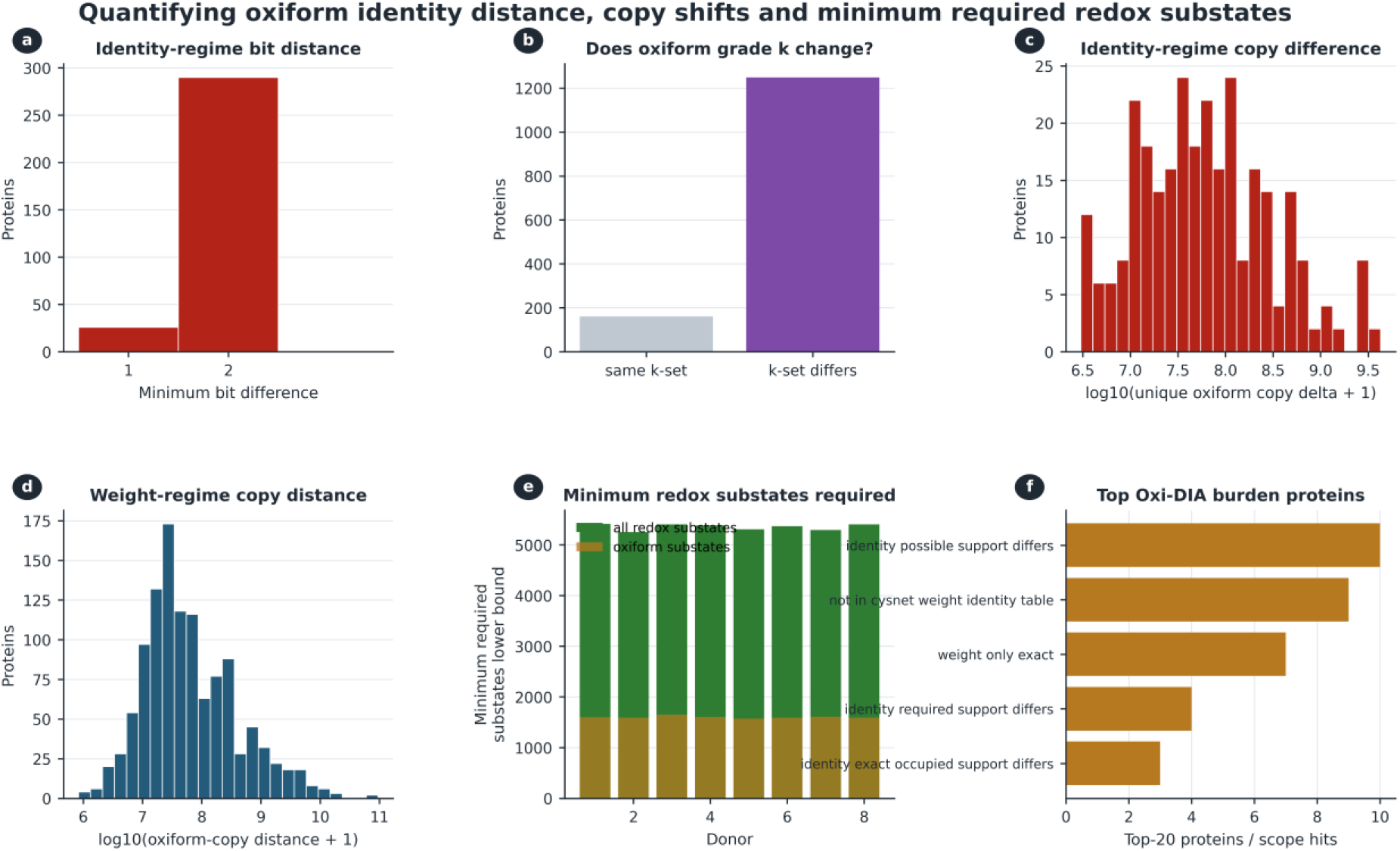
Quantifying oxiform identity distance, copy-number shifts and minimum redox substate burden. **(A)** Minimum bit-distance between donor-specific identity-regime oxiform states, showing that most identity differences require a two-coordinate cysteine-redox change rather than a single-site change. **(B)** Classification of whether identity-regime differences preserve the same oxidation grade *k* or alter the occupied *k*-set, indicating that most donor differences involve changes in the number of oxidised cysteine coordinates represented in the compatible oxiform support. **(C)** Distribution of copy-number differences for identity-regime oxiforms, expressed as log_10_(unique oxiform copy delta + 1). **(D)** Distribution of copy-number distances for weight-regime differences, expressed as log_10_(oxiform-copy distance + 1), capturing cases where the same oxiform identity is retained but its abundance differs between donors. **(E)** Minimum required number of redox substates per donor, partitioned into all redox substates and oxiform substates, showing the lower-bound substate burden implied by the measured cysteine-redox marginals. **(F)** Proteins contributing most frequently to the top Oxi-DIA burden across identity and weight regimes, highlighting recurrent proteins that dominate donor-level oxiform variation.

Having defined the empirical input regime, we next applied the CysNet theorem engine to determine which oxiform-level constraints were implied from the Oxi-DIA data. Across the eight donor cell lines, CysNet analysed 49,940 protein-donor records, of which only 2,820 had complete cysteine coverage while 47,120 were incompletely covered. Hence, the empirical, bottom-up redox proteomics was dominated by incomplete protein records^17^, a fact that has led to the characterisation of oxiform distributions being considered as impossible using bottom-up MS data. Nonetheless, constraint-based inference of the oxiform distributions should remain informative.

Among complete-coverage records, CysNet identified 2,256 exact singleton feasible oxiform ensembles and 564 bounded non-singleton feasible oxiform ensembles. Hence, approximately 80% of complete-coverage records collapsed to an exact ensemble distribution, whereas the remaining complete records retained ambiguity, but were bounded. In incomplete records, CysNet returned existence and exclusion constraints. Across the cohort, 15,441 incomplete protein-donor records contained at least one positive measured cysteine marginal and therefore required occupancy of at least one oxidised state within the oxiform state space. Including complete records, 16,763 protein-donor records were mathematically consistent with the existence of an oxiform with at least one oxidised cysteine residue^8^.

Site at the tropical boundary—fully reduced or oxidised—produced extensive oxiform state-space collapse. Across the cohort, boundary sites (e.g., fully reduced = 0) removed 126,879 binary dimensions of unconstrained state space. Every time a fully reduced or oxidised site was observed, the state space collapsed dramatically, often by orders of magnitude, limiting the number of unique oxiforms that could exist in the sample. A further constraint arose from partial redox values, which bounded by mathematical constraint the potential occupancy of thousands of oxiforms.

Next, CysNet scaled these constraints to protein copy number. Across 49,628 records with copy-number estimates, the measured cysteine-containing proteome comprised 4.40 × 10^13^ protein copies. CysNet bounded the cohort oxiform content at 6.36 × 10^12^–8.24 × 10^12^ copies, corresponding to 14.44–18.71% of the measured copy pool. Fully reduced-compatible oxiforms accounted for 3.58 × 10^13^–3.77 × 10^13^ copies, or 81.29–85.56% of the measured pool. Hence, from the bottom-up redox proteomic data, CysNet inferred copy-number scaled oxiform differences between the different cell lines—a feat previously considerd intractable on bottom-up data by its construction.

### CysNet decomposes redox variation into oxiform weight and identity regimes

Conceptually, redox variation between biological samples could occur either through changes in oxiform identity (composition) and/or weight (intensity). In the identity regime, samples could differ in which oxiforms are compatible or occupied. For example, 010 in one sample compared to 001 in another. This represents a compositional change in the oxiform ensemble, placing proteoform heterogeneity itself as a mechanism of redox regulation. In the weight regime, the same oxiforms are retained, but their fractional occupancies differ across the population of molecules in the respective protein group. For example, 75% versus 50% occupancy of the same state in different samples. This represents an intensity change in the oxiform ensemble, placing the extent of oxidation within a conserved oxiform identity structure as the regulatory variable. By deriving oxiform-level inferences, CysNet can quantify the relative contribution of these regimes to biological redox variation.

We therefore classified cell line-associated oxiform variation across a hierarchy of evidence levels. In the strongest class, complete-coverage proteins with exact solutions in all donors, CysNet resolved 43 multi-cysteine proteins into donor cell line-specific oxiform ensembles. Of these, 28 proteins, (65.1%), showed weight-only variation, in which the same oxiform identities were present in every line but their fractional occupancies differed. Exactly 15 proteins, (34.9%), showed donor-dependent changes in the exact occupied oxiform identities. When all complete-coverage proteins were considered, including bounded non-singleton solutions, CysNet classified 115 proteins: 52 (45.2%), preserved the same oxiform identity structure but differed in weights or bounds, whereas 63 (54.8%), showed identity or feasibility differences through changes in required, possible or exactly occupied substates. Hence, both regimes contributed to biological redox variation,

Next, we analysed cases whereby Oxi-DIA did not detect all of the cysteine residues belonging to the detected protein group (i.e., the incomplete cysteine coverage set). In this regard, 565 of 1,262 proteins (44.8%) showed same-identity weight or bound variation and 697 (55.2%) showed oxiform identity or feasibility variation. Even when the same site was detected in these proteins, 513 of 1,147 proteins (44.7%) showed same-identity variation, whereas 634 (55.3%) showed altered oxiform identity or feasibility. Hence, donor cell line-associated redox variation was explained by both differences in the extent of oxidation at some cysteine sites, as well as by differences in which cysteine sites were oxidised.

We next quantified the contributions of each regime between the respective cell lines. For identity-regime classifications, we asked how far donor cell line-specific oxiform identities moved in binary state space. Across 1,409 identity-regime protein-scope entries, the minimum donor-to-donor bit distance ranged from one to two cysteine-coordinate changes, with a median minimum distance of two. Moreover, 1,249 entries (88.6%) changed their accessible oxidation-grade structure. This means that the potential to access oxiforms with more oxidised cysteine residues differed. As a result, we observed cell line differences in the number of oxidised cysteines that needed to explain bottom-up redox proteomic data.

Copy-number scaling showed that these identity shifts were substantial in terms of the number of molecules involved: the median unique oxiform copy-number difference associated with identity-specific substates was (5.41 × 10^7^) copies. Variation in redox weight at the same cysteine sites affected a similarly large number of molecules in the respective proteomes. Across 1,158 same-identity protein-scope entries, the median oxiform-copy distance was (4.53 × 10^7^) copies, with the largest same-identity shift reaching (5.41 × 10^10^) copies. Hence, both cysteine site identity and redox weight regimes were quantitatively important contributors to the observed proteomic variation in oxiforms between the different cell lines.

Finally, we asked how many oxiforms, were minimally required to explain the measured data. Within the dataset, CysNet inferred donor cell line-specific lower bounds of 5,253–5,412 total oxiform states and 1,571–1,650 states with at least one oxidised cysteine per donor. These are conservative lower bounds: they count only the minimum number of observed-coordinate states required by the measured redox values at each cysteine site and do not assign unmeasured cysteine coordinates when the site was not detected. Even under this conservative interpretation, the donor cell line-specific iPSC biological redox variation was inferentially mapped back to changes in the identity (compositions) and weight (abundance) of thousands of oxiforms.

### CysNet is provided as an open-source theorem-constrained Python workflow

CysNet is provided as an open-source, theorem-constrained Python workflow for inferring oxiform constraints from bottom-up cysteine redox proteomics data. The software takes reduced and reversibly oxidised cysteine site matrices, the FASTA file used for the proteomic search, and optional protein-group abundance values as input. It then calculates residue-resolved cysteine redox value, maps the detected cysteine sites onto FASTA-derived protein group, classifies complete and incomplete cysteine coverage at the protein group level, applies the oxiform constraint theorem, and returns protein-level oxiform constraint reports.

CysNet is distributed with multiple access routes. Command-line tools are provided for theorem examples, Oxi-DIA light/heavy site import and FASTA cysteine site mapping, while the full constraint and copy-number workflows can be run through the Python API. For non-command-line use, CysNet includes a Colab-native upload workflow and a desktop graphical interface that runs the same tested pipeline. The graphical interface provides a full pipeline mode for processing experimental datasets and a theorem explorer for inspecting how cysteine marginals constrain individual oxiform state spaces.

Hence, CysNet can be immediately implemented both on new (as here) and archived (e.g., OxiMouse) redox proteomic datasets without the need to know how to code.

## Discussion

CysNet is a computational workflow that is designed to infer additional information from conventional bottom-up redox proteomic data. By implementing a theorem-constrained workflow, CysNet establishes an oxiform inference layer for bottom-up redox proteomics. Thus, CysNet converts cysteine oxidation measurements from isolated, site-level percentages into exact, mathematical constraints on the underlying ensemble of oxiforms. This effectively changes the analysis object: each cysteine site is no longer interpreted alone, but instead as a part of an oxiform state space. When applied here to analyse redox proteomes in distinct human iPS cell lines, this revealed biological redox variation. Further, the observed donor cell line-associated redox variation could be attributed into variation arising from both differences in the extent of oxidation at the same cysteine sites (oxiform weight) as well as differences in which specific cysteine residues were oxidised (oxiform identity). CysNet recovered proteoform-level structure that was latent in the original bottom-up MS data.

Conceptually, CysNet treats bottom-up MS redox measurements of cysteine oxidation as marginals related to a hidden oxiform ensemble that defines the proteome in the biological sample being analysed, rather than as independent, site-level endpoints. Using bottom-up MS redox proteomic datasets as input, CysNet returns the exact constraint class supported by the data: compatible oxiform substates, excluded substates, required substates, exact solutions and bounded feasible oxiform ensembles. For example, in a protein with three cysteine residues, cysteine oxidation values of 0, 0, 0.25 can only belong to an oxiform distribution of 000 (75% of molecules with all cysteines fully reduced) and 001 (25% of molecules with the third site cysteine oxidised and the other two cysteines reduced) by theorem. Note that theorem still mathematically constrains the feasible oxiform state space, even if only one site is measured, ensuring every cysteine site whose redox state is measured in the bottom-up MS data also contributes information. Hence, CysNet introduces an oxiform state space inference layer into redox proteomics data.

Using CysNet to analyse oxiform distributions in eight different human iPSC redox proteomes form genetically distinct donors revealed proteoform-level structure that was latent in the original Oxi-DIA MS measurements. Analysis of the redox state of different cysteine sites established donor cell line-associated cysteine-redox variation and copy-number scaling showed that these differences between the respective cell lines corresponded to a substantial change in the oxidised molecular burden. CysNet resolved how this cysteine redox variation was represented in oxiform state space. Across evidence tiers, donor variation separated into two oxiform regimes: many proteins preserved the same set of compatible oxiform identities but changed the fraction of molecules oxidised at the same cysteine sites, whereas other proteins changed which oxiforms were represented in the expressed proteome ensemble, with some cysteine sites oxidised in a cell line-specific pattern.

Quantitatively, redox identity differences often involved two cysteine-coordinate changes and frequently altered oxidation grade (*k*), while both identity and weight regimes carried tens of millions of copy-number-scaled oxiform differences. These data represent a significant advance in the ability to characterise expressed proteomes that has been enabled by CysNet: here donor cell line-associated redox variation could be identified and further revealed to result from differences between the cell lines in both oxiform occupancy and changes in oxiform identity. This establishes that redox variation depends on both the magnitude of cysteine oxidation (how many molecules in each protein group are oxidised or reduced at each cysteine site) and which combinations of cysteine residues at different sites in a molecule are reduced or oxidised, respectively. Hence, in the comparison of human iPS cell lines from different female donors, cysteine redox variation is defined by changes in the composition, occupancy and accessible regions of the proteoform state space.

While here CysNet has specifically addressed latent information in bottom-up MS data sets concerning proteoform variation at the level of cysteine oxidation the broader implication is that many MS datasets relating to other post translational protein modifications (PTMs) may also contain valuable latent proteoform-level information. CysNet provides one implementation for cysteine, but the underlying mathematics is general. Future extensions of CysNet will therefore adapt the same logic to explore proteoform variation in other PTMs, such as phosphorylation to constrain phosphoforms (i.e., PhospoNet). We propose this can help to advance proteomics by maximising the information contained in bottom-up MS measurements to quantify modified sites and then to infer the proteoform state spaces those sites constrain.

## Methods

### Human induced pluripotent stem cells

Eight, female donor-derived human iPSC lines used were cultured as previously described ^22,32^. Briefly, the human iPSC lines were cultured in Essential 8 (E8) medium (E8 complete medium supplemented with (50x) E8 supplement ThermoFisher-A1517001) on tissue-culture dishes coated with 10 μg/cm2 of reduced Growth Factor Basement Membrane matrix (Geltrex, ThermoFisher A1413202 resuspended in basal medium DMEM/F12 Thermo Fisher 21331020). Cells were routinely tested for mycoplasma (MycoStrip, InvivoGen, UK). Approximately 2×10^6^ cells/ml were collected per biological replicate. Each female had three biological replicates, yielding 24 discrete samples. To avoid storage-induced oxidation the cells were processed immediately (see Cysteine labelling).

### Cysteine labelling

To label cysteines for Oxi-DIA, samples were lysed in fresh light-protected lysis buffer (2%SDS [ThermoFisher, UK, #15553027), 100 mM HEPES (pH 7.4[Sigma, UK, #0000338713), 2 mM EDTA [Sigma, UK, #03690], 1 mM DPTA [Sigma, UK, #D6518], and 0.1 mM neocuprine[Sigma, UK, #121908]) supplemented with 25 mM light *N*-ethylmaleimide (NEM_L, Sigma, UK, #04260). The NEN_L was added just before use to prevent aqueous hydrolysis to cysteine unreactive maleimic acid^33^. This method has been shown to minimise lysis-induced oxidation^24^. After clarifying the lysate by centrifugation (5,000 *g* for 2 min), the samples were incubated for 1 h in the dark at 37℃ to enable NEM-L to alkylate reduced cysteines via a covalent thioether bond. Per our previous work^24^, sonication and boiling were omitted to preserve the endogenous cysteine redox states.

After completing the NEM_L labelling reaction, the sample was cleaned up via an organic solvent-based extraction procedure ^34^. The pellets were subsequently resuspended in lysis buffer supplemented with 10 mM of neutral-TCEP (ThermoFisher, UK, #77720) to reduce reversibly oxidised cysteines via nucleophilic phosphine chemistry^35^. After an incubation period of 20 min, 15 mM heavy *N*-ethyl-d_5_-maleimide (NEM_H, Sigma, UK, #692905) was added to the lysate to alkylate newly reduced (oxidised) cysteines^36^. After completing the differential alkylation reactions, the samples were cleaned via organic phase extraction.

The pellets were resuspended in 50 mM ammonium bicarbonate and digested overnight at 37℃ in the dark using 1-10 µg sequence-grade Trypsin and LysC (ThermoFisher, UK, #A40009). The digestion reaction was quenched using 1% FA. The volatile ABC salt was removed via drying the peptides at 30℃ to dryness in a speedvac. Peptides were resuspended in 1% DMSO and 0.1%FA. Peptide concentration was determined using a fluorescent microplate-based assay (ThermoFisher, UK, #23290).

### Oxi-DIA

To measure the cysteine redox state of the sample via Oxi-DIA, 500 nanogram of peptide input was loaded onto a Vanquish Neo U-HPLC coupled to an Orbitrap Astral™ mass spectrometer^25,26^, which was mass and system calibrated in positive ion mode. After passing through a 5 cm trap column, the samples were loaded onto a 25 cm IonOptics C_18_ column and resolved over 42-min with the ion source set to between 1600-2500 V. Mobile phase A and B were 0.1%-FA and 80% acetonitrile (ACN) in 0.1%-FA, respectively. By adjusting the mobile phase, the peptides were resolved using a nonlinear gradient 5-80% ACN. MS^1^ (intact peptides) spectra were acquired in the Orbitrap every 0.6 seconds at the full instrument resolving power of 240,000. After fragmenting peptides via high-energy collisional induced dissociation at 25%-energy, MS^2^ spectra were collected in the Astral across the 380-980 m/z range at full instrument resolving power, automatic gain control of 500%, and a maximal injection time of 3 milliseconds. The 380-980 m/z range was split into 300, 2-Thompson windows for narrow DIA ^29^. In this work, each biological replicate (*n* = 24) was injected three times, yielding 72 raw files. Hence, 9 raw files belonged to each donor.

### DIA-NN

The Thermo raw files were analysed in <u>DIA-NN</u> (v.2.2)^27^ using an *in silico* predicted spectral library generated from a species-specific FASTA file. The spectral library was generated with the following instructions inputted into the command line interface: “-*-var-mod UniMod:108,125*.*047,C*”, “*--var-mod UniMod:776, 130*.*076,C*”. In addition, variable N-terminal methionine excision and up to 1 missed tryptic digestion were allowed. The maximum number of variable modifications per peptide was set to 3 and the peptide length range considered was 5-35 amino acids. For the quantitative search, the raw files were searched against the *in silico* generated spectral library. For this search, “*--export quant*” was added to the command line interface. The other search settings remained the same. Match between runs (MBR) and peptidoform scoring mode were enabled. In addition, quant-UMS (precision) was selected to minimise ratio distortion^28^. Peptide and protein false discovery rate (FDR) was set to 0.01 and cysteine residues with site localisation probabilities of 0.9 were computationally analysed^37^.

### Redox analysis

To compute the site-wise cysteine oxidation per sample and then scale this to protein groups, the relevant DIA-NN output files were processed using custom Python scripts implemented in Jupyter notebooks and executed within a high RAM Google Colab runtime^8^. To do so, adapted scripts from our previous work (https://github.com/JamesCobley/ReCap), were implemented. The CysNet theorem detailed in the supplement, was computationally instantiated using the CysNet python package in a google colab environment. All of the details, including each individual script, of the implementation are available at https://github.com/JamesCobley/CysNet/tree/main.

