## Supplemental Information for "CysNet: Theorem constrained inference of cysteine redox proteoform states from bottom-up mass spectrometry data"

**Supplemental Note 1. Feasible cysteine proteoform distributions form a convex polytope**

Let

$$\Omega_{r}=\{0,1{\}}^{r}$$

denote the binary cysteine proteoform state space for a protein with $r$cysteine residues, where $0$denotes reduced and $1$denotes oxidised. Let

$$m=(m_{1},\ldots,m_{r})\in[0,1]^{r}$$

be the vector of measured site-wise cysteine redox marginals. A proteoform distribution is a vector

$$p=(p_{x})_{x\in\Omega_{r}}\in\mathbb{R}^{2^{r}}$$

satisfying

$$p_{x}\geq0\forall x\in\Omega_{r},$$

and

$$\sum_{x\in\Omega_{r}} p_{x}=1.$$

The set of all proteoform distributions consistent with the measured marginals is

$$P(m)=\left\{ p\in\mathbb{R}^{2^{r}}:p_{x}\geq0,\text{ }\sum_{x\in\Omega_{r}} p_{x}=1,\text{ }\sum_{x\in\Omega_{r}} x_{i}p_{x}=m_{i}\text{ for all }i=1,\ldots,r \right\}.$$

For the binary cysteine state space, every vector $m\in[0,1]^{r}$is feasible. Indeed, feasibility is witnessed by the independent product distribution

$$p_{x}=\prod_{i=1}^{r} m_{i}^{x_{i}}(1-m_{i})^{1-x_{i}}.$$

This distribution has site-wise marginals $m_{i}$, so $P(m)\neq\emptyset$for every $m\in[0,1]^{r}$.

**Theorem 1. Convex polytope structure**

For any marginal vector $m\in[0,1]^{r}$, the feasible set $P(m)$is a non-empty convex polytope.

**Proof**

The probability simplex in $\mathbb{R}^{2^{r}}$is

$$\Delta_{2^{r}-1}=\left\{ p\in\mathbb{R}^{2^{r}}:p_{x}\geq0\text{ }\forall x,\text{ }\sum_{x\in\Omega_{r}} p_{x}=1 \right\}.$$

This is a convex polytope, since it is defined by finitely many linear inequalities and one linear equality. For each residue $i$, the marginal constraint

$$\sum_{x\in\Omega_{r}} x_{i}p_{x}=m_{i}$$

defines an affine hyperplane. Therefore,

$$P(m)=\Delta_{2^{r}-1}\cap\bigcap_{i=1}^{r} \left\{ p\in\mathbb{R}^{2^{r}}:\sum_{x\in\Omega_{r}} x_{i}p_{x}=m_{i} \right\}.$$

Thus $P(m)$is the intersection of a convex polytope with finitely many affine hyperplanes. Hence $P(m)$is a convex polytope. Non-emptiness follows from the product-distribution construction above. ∎

**Corollary 1. Deterministic marginals impose hard exclusion faces**

If $m_{i}=0$for some residue $i$, then every feasible distribution $p\in P(m)$assigns zero probability to all proteoforms with $x_{i}=1$. Similarly, if $m_{i}=1$, then every feasible distribution assigns zero probability to all proteoforms with $x_{i}=0$.

**Proof**

Suppose $m_{i}=0$. Then

$$\sum_{x\in\Omega_{r}} x_{i}p_{x}=0.$$

Since $x_{i}p_{x}\geq0$for every $x$, every term with $x_{i}=1$must satisfy $p_{x}=0$. Therefore, all proteoforms containing oxidation at residue $i$are excluded.

The case $m_{i}=1$is analogous. Since

$$\sum_{x\in\Omega_{r}} x_{i}p_{x}=1$$

and

$$\sum_{x\in\Omega_{r}} p_{x}=1,$$

all probability mass must lie on states with $x_{i}=1$. Hence all proteoforms with $x_{i}=0$are excluded. ∎

**Corollary 2. Singleton solutions after deterministic exclusions**

Let

$$I=\{i:m_{i}\in(0,1)\}$$

be the set of residues with intermediate, non-deterministic marginals. After applying the hard exclusions induced by residues with $m_{i}\in\{0,1\}$, the feasible distribution is supported only on proteoforms that agree with the deterministic residues. Therefore, the remaining admissible state space contains at most

$$2^{\mid I\mid}$$

proteoforms.

If $\mid I\mid=0$, then $P(m)$is a singleton. All residues are deterministic, so exactly one proteoform is compatible with the measured marginals.

If $\mid I\mid=1$, then $P(m)$is also a singleton. In this case, all residues except one are deterministic. Let $j$be the single residue with $m_{j}\in(0,1)$. After hard exclusions, only two proteoforms remain possible: one with $x_{j}=0$and one with $x_{j}=1$. Writing these states as $x^{\left( 0 \right)}$and $x^{\left( 1 \right)}$, the marginal constraint gives

$$p_{x^{\left( 1 \right)}}=m_{j},$$

and normalization gives

$$p_{x^{\left( 0 \right)}}=1-m_{j}.$$

Thus the feasible distribution is uniquely determined.

If $\mid I\mid\geq2$, site-wise marginals alone generally do not uniquely determine the joint proteoform distribution, because correlations among the intermediate residues remain unspecified. After deterministic exclusions, the feasible set lies in an affine subspace of dimension at most

$$2^{\mid I\mid}-\mid I\mid-1,$$

corresponding to $2^{\mid I\mid}$possible remaining proteoforms, one normalization constraint, and $\mid I\mid$marginal constraints. Therefore, for $\mid I\mid\geq2$, the feasible set typically has positive dimension.

In particular, the independent product distribution

$$p_{x}=\prod_{i\in I} m_{i}^{x_{i}}(1-m_{i})^{1-x_{i}}$$

is always one feasible point on the reduced state space, and it is the unique maximum-entropy solution under the marginal constraints. However, unless additional information specifies independence or higher-order correlations, it is not generally the unique feasible proteoform distribution.

**Example**

For $r=3$, let the measured marginals be

$$m=(0,0,0.25).$$

The first two residues have deterministic marginals. By Corollary 1, all proteoforms with oxidation at residue 1 or residue 2 are excluded. Therefore, the only compatible states are

$$000\text{and}001.$$

The third marginal requires

$$p_{001}=0.25.$$

Normalization then gives

$$p_{000}=0.75.$$

Hence the feasible set collapses to the unique cysteine proteoform distribution

$$p_{000}=0.75,p_{001}=0.25.$$

In this case, site-resolved peptide measurements do not merely constrain the proteoform ensemble; they uniquely reconstruct it.

**Interpretation**

Site-resolved peptide measurements do not generally specify a single cysteine proteoform distribution. Instead, they define a feasible region $P(m)$inside the full proteoform simplex. This feasible region contains all joint proteoform distributions whose residue-wise oxidation marginals agree with the measured peptide-level data.

Boundary marginals have a special role. Exact values of $0$or $1$act as hard exclusion rules, eliminating all proteoforms incompatible with the deterministic residue state. In a protein with many cysteine residues, even a single deterministic site can remove an exponentially large subset of the binary proteoform state space.

Intermediate marginals impose linear constraints but do not, by themselves, specify correlations between residues. Therefore, when multiple cysteine residues remain partially oxidised, the measured marginals define a family of compatible proteoform distributions rather than a unique solution. However, when deterministic marginals reduce the remaining unresolved space to at most one binary coordinate, the feasible polytope collapses to a single point. In that regime, bottom-up site-resolved redox measurements uniquely determine the cysteine proteoform distribution.

This provides the formal basis for using site-wise cysteine redox measurements not only to constrain proteoform state space, but, in defined cases, to solve the proteoform distribution exactly.

**Supplementary Note 2. OxiTope imposes chemical partition constraints on the cysteine proteoform polytope**

**1. Proteoform state space**

For a protein with $r$cysteine residues, define the binary cysteine proteoform state space as

$$\Omega_{r}=\{0,1{\}}^{r},$$

where $0$denotes reduced and $1$denotes reversibly oxidised. A proteoform distribution is

$$p=(p_{x})_{x\in\Omega_{r}},$$

with

$$p_{x}\geq0,\sum_{x\in\Omega_{r}} p_{x}=1.$$

The all-reduced proteoform is denoted

$$0_{r}=(0,0,\ldots,0).$$

The oxiform subspace is the complement

$$\Omega_{r}^{\mathrm{ox}}=\Omega_{r}\setminus\{0_{r}\}.$$

Thus,

$$\Omega_{r}=\{0_{r}\}\cup\Omega_{r}^{\mathrm{ox}}.$$

**2. OxiTope chemical partition**

OxiTope introduces a binary chemical gate that partitions intact protein molecules into two experimentally observable fractions:

$$F_{\mathrm{red}}$$

and

$$F_{\mathrm{ox}}.$$

Under ideal gate performance,

$$F_{\mathrm{red}}\leftrightarrow\{0_{r}\}$$

and

$$F_{\mathrm{ox}}\leftrightarrow\Omega_{r}^{\mathrm{ox}}.$$

That is, molecules in the reduced fraction occupy the all-reduced vertex, whereas molecules in the oxiform fraction occupy the complementary subspace containing at least one reversibly oxidised cysteine.

Let

$$q_{\mathrm{red}}$$

and

$$q_{\mathrm{ox}}$$

denote the experimentally measured fraction of protein signal assigned to the reduced and oxiform fractions, respectively, after yield-scaled protein quantification.

Then

$$q_{\mathrm{red}}+q_{\mathrm{ox}}=1.$$

The OxiTope gate imposes the linear constraints

$$p_{0_{r}}=q_{\mathrm{red}},$$

and

$$\sum_{x\in\Omega_{r}^{\mathrm{ox}}} p_{x}=q_{\mathrm{ox}}.$$

Since the second constraint follows from normalization and the first, the essential new constraint is

$$p_{0_{r}}=q_{\mathrm{red}}.$$

**Theorem 1. OxiTope constrains the proteoform simplex by fixing the all-reduced vertex mass**

Let

$$\Delta_{2^{r}-1}=\left\{ p\in\mathbb{R}^{2^{r}}:p_{x}\geq0,\text{ }\sum_{x\in\Omega_{r}} p_{x}=1 \right\}$$

be the cysteine proteoform probability simplex. Under ideal OxiTope partitioning, the feasible set of proteoform distributions consistent with the measured reduced fraction $q_{\mathrm{red}}$is

$$P_{\mathrm{OxiTope}}(q_{\mathrm{red}})=\left\{ p\in\Delta_{2^{r}-1}:p_{0_{r}}=q_{\mathrm{red}} \right\}.$$

Then $P_{\mathrm{OxiTope}}(q_{\mathrm{red}})$is a convex polytope. If $q_{\mathrm{red}}=1$, the feasible set collapses to the singleton all-reduced distribution. If $q_{\mathrm{red}}=0$, the all-reduced vertex is excluded.

**Proof**

The probability simplex $\Delta_{2^{r}-1}$is a convex polytope. The constraint

$$p_{0_{r}}=q_{\mathrm{red}}$$

is an affine hyperplane. Therefore,

$$P_{\mathrm{OxiTope}}(q_{\mathrm{red}})=\Delta_{2^{r}-1}\cap\{p:p_{0_{r}}=q_{\mathrm{red}}\}$$

is the intersection of a convex polytope with an affine hyperplane, and is therefore a convex polytope.

If $q_{\mathrm{red}}=1$, normalization requires

$$p_{0_{r}}=1$$

and

$$p_{x}=0\text{for all}x\neq0_{r}.$$

Thus the feasible set is the singleton all-reduced distribution.

If $q_{\mathrm{red}}=0$, then

$$p_{0_{r}}=0,$$

so the all-reduced vertex is excluded and all probability mass lies in the oxiform subspace. ∎

**3. Incorporating residue-level Oxi-DIA marginals**

Oxi-DIA provides site-wise oxidation marginals

$$m_{i}=\sum_{x\in\Omega_{r}} x_{i}p_{x},i=1,\ldots,r.$$

Combining OxiTope with Oxi-DIA gives the feasible set

$$P(q_{\mathrm{red}},m)=\left\{ p\in\Delta_{2^{r}-1}:p_{0_{r}}=q_{\mathrm{red}},\text{ }\sum_{x\in\Omega_{r}} x_{i}p_{x}=m_{i}\text{ for all }i=1,\ldots,r \right\}.$$

This is the central mathematical object of OxiTope-CysNet inference.

**Theorem 2. OxiTope-CysNet inference is a convex polytope problem**

For any experimentally compatible $q_{\mathrm{red}}\in[0,1]$and marginal vector $m\in[0,1]^{r}$, the feasible set

$$P(q_{\mathrm{red}},m)$$

is a convex polytope.

**Proof**

The feasible set is obtained by intersecting the probability simplex with finitely many affine hyperplanes:

$$p_{0_{r}}=q_{\mathrm{red}}$$

and

$$\sum_{x\in\Omega_{r}} x_{i}p_{x}=m_{i},i=1,\ldots,r.$$

An intersection of a convex polytope with finitely many affine hyperplanes is a convex polytope. ∎

**4. Copy-number-weighted occupancy**

Let $N$denote the estimated copy number of the protein. Define the copy-number-weighted occupancy of proteoform $x$as

$$w_{x}=Np_{x}.$$

Then

$$w_{x}\geq0,\sum_{x\in\Omega_{r}} w_{x}=N.$$

OxiTope fixes the copy number assigned to the all-reduced vertex:

$$N_{\mathrm{red}}=Np_{0_{r}}=Nq_{\mathrm{red}},$$

and the copy number assigned to the oxiform subspace:

$$N_{\mathrm{ox}}=Nq_{\mathrm{ox}}=N(1-q_{\mathrm{red}}).$$

Thus,

$$\sum_{x\in\Omega_{r}^{\mathrm{ox}}} w_{x}=N_{\mathrm{ox}}.$$

This remains true even when the internal distribution within $\Omega_{r}^{\mathrm{ox}}$is unresolved.

**Corollary 1. OxiTope fixes oxiform copy number even when oxiform identity is unresolved**

Under ideal OxiTope partitioning, the total number of molecules occupying the oxiform subspace is

$$N_{\mathrm{ox}}=N(1-q_{\mathrm{red}}).$$

Therefore, even if the individual oxiform states $x\in\Omega_{r}^{\mathrm{ox}}$are not uniquely resolved, their total molecular occupancy is fixed.

**Interpretation**

OxiTope does not require complete residue-level resolution to quantify the oxiform population. The internal distribution over oxiform states may remain a polytope, but the total probability mass and copy-number occupancy of that subspace are experimentally constrained.

**5. Copy-number bounds on realised state occupancy**

The theoretical number of cysteine proteoforms is

$$2^{r}.$$

However, if a protein has only $N$copies, then no more than $N$proteoform states can be materially occupied:

$$\mid supp(p)\mid\leq\min(2^{r},N),$$

assuming discrete molecule counts.

For the oxiform subspace,

$$\mid\mathrm{supp}_{\mathrm{ox}}(p)\mid\leq\min(2^{r}-1,N_{\mathrm{ox}}).$$

Thus, if

$$N_{\mathrm{ox}}\ll2^{r}-1,$$

most theoretically possible oxiform states cannot be occupied by real molecules.

**Corollary 2. OxiTope converts theoretical state space into weighted occupancy geometry**

OxiTope transforms cysteine proteoform inference from an unweighted enumeration of possible states into a copy-number-weighted occupancy problem. The theoretical oxiform subspace may contain

$$2^{r}-1$$

states, but the realised oxiform ensemble is bounded by

$$N_{\mathrm{ox}}.$$

Therefore, the relevant biological question is not only which oxiforms are mathematically possible, but how a finite number of protein copies is distributed across the chemically constrained oxiform subspace.

**6. Monothiol proteins**

For a monothiol protein,

$$r=1,$$

so

$$\Omega_{1}=\{0,1\}.$$

The all-reduced vertex is

$$0_{1}=0,$$

and the oxiform subspace is

$$\Omega_{1}^{\mathrm{ox}}=\{1\}.$$

Therefore,

$$p_{0}=q_{\mathrm{red}},$$

and

$$p_{1}=q_{\mathrm{ox}}=1-q_{\mathrm{red}}.$$

**Corollary 3. For monothiol proteins, OxiTope directly reports the residue redox state**

For a protein containing a single cysteine residue, the OxiTope reduced/oxiform partition uniquely determines the cysteine redox distribution:

$$p_{\mathrm{reduced}}=q_{\mathrm{red}},$$

$$p_{\mathrm{oxidised}}=q_{\mathrm{ox}}.$$

Therefore,

$$\%Ox=100q_{\mathrm{ox}}.$$

This holds even if the cysteine-containing peptide itself is not detected, provided the protein-level fraction assignment is measured.

**Proof**

For $r=1$, the oxiform condition “at least one reversibly oxidised cysteine” is equivalent to “the only cysteine is oxidised.” Therefore, the OxiTope fraction split is identical to the binary residue redox split. ∎

**7. Experimental validation conditions**

The formal inference above assumes ideal gate performance. Experimentally, this requires validation that:

$$F_{\mathrm{red}}$$

contains no detectable residual reversibly oxidised cysteine signal, and that

$$F_{\mathrm{ox}}$$

is chemically specific for cysteine-dependent SPDP capture.

The essential controls are:

1. **Reduced-channel breakthrough control**
   After SPDP capture, the reduced fraction is challenged with TCEP and fluorescent maleimide. Signal at background indicates no detectable residual reducible cysteine pool.
2. **Oxiform-channel positivity control**
   The bead-eluted fraction should contain reducible cysteine signal.
3. **Cysteine-free protein exclusion control**
   Proteins lacking cysteine residues should be absent or background-level in the TCEP-eluted oxiform fraction. Such proteins may passively associate with beads, but have no chemical route into the SPDP-linked oxiform pool and should not be released by reductive elution after stringent washing.

These controls define the conditions under which the chemical partition can be interpreted as a valid constraint on the proteoform polytope.

**8. Summary**

CysNet begins with site-wise residue marginals and defines a feasible proteoform polytope. OxiTope adds an upstream chemical partition that fixes the probability mass of the all-reduced vertex and the complementary oxiform subspace. This converts incomplete residue-level evidence into a constrained, copy-number-weighted occupancy problem. In complete-coverage cases, residue-level CysNet can resolve the internal oxiform structure. In incomplete-coverage cases, OxiTope still fixes the total oxiform probability and copy number, thereby placing a strong experimental constraint on what proteoform distributions can exist.
